# Quantifying and correcting slide-to-slide variation in multiplexed immunofluorescence images

**DOI:** 10.1101/2021.07.16.452359

**Authors:** C.R. Harris, E.T. McKinley, J.T. Roland, Q. Liu, M.J. Shrubsole, K.S. Lau, R.J. Coffey, J. Wrobel, S.N. Vandekar

**Affiliations:** Department of Biostatistics, Vanderbilt University Medical Center, Nashville, TN, USA; Epithelial Biology Center, Vanderbilt University Medical Center, Nashville, TN, USA; Department of Cell and Developmental Biology, Vanderbilt University School of Medicine, Nashville, TN, USA; Department of Surgery, Vanderbilt University School of Medicine, Nashville, TN, USA; Center for Quantitative Sciences, Vanderbilt University Medical Center, Nashville, TN, USA; Division of Epidemiology, Vanderbilt Ingram Cancer Center, Vanderbilt University Medical Center, Nashville, TN, USA; Division of Gastroenterology, Hepatology, and Nutrition, Department of Medicine, Vanderbilt University Medical Center, Nashville, TN, USA; Department of Biostatistics & Informatics, Colorado School of Public Health, Aurora, CO, USA

## Abstract

**Motivation:** The multiplexed imaging domain is a nascent single-cell analysis field with a complex data structure susceptible to technical variability that disrupts inference. These in situ methods are valuable in understanding cell-cell interactions, but few standardized processing steps or normalization techniques of multiplexed imaging data are available.

**Results:** We implement and compare data transformations and normalization algorithms in multiplexed imaging data. Our methods adapt the ComBat and functional data registration methods to remove slide effects in this domain, and we present an evaluation framework to compare the proposed approaches. We present clear slide-to-slide variation in the raw, unadjusted data, and show that many of the proposed normalization methods reduce this variation while preserving and improving the biological signal. Further, we find that dividing this data by its slide mean, and the functional data registration methods, perform the best under our proposed evaluation framework. In summary, this approach provides a foundation for better data quality and evaluation criteria in the multiplexed domain.

**Availability and Implementation:** Source code is provided at https://github.com/statimagcoll/MultiplexedNormalization.

**Supplementary information:** Supplementary information is available online.

## Introduction

Single-cell assays are increasingly valued for their ability to provide information about the cell micro-environment and cell population interactions in healthy and cancerous tissues (Islam et al., 2020; McKinley et al., 2019; Shrubsole et al., 2008). Multiplexed imaging methods like multiplexed immunofluorescence (MxIF) (Gerdes et al., 2013), multiplexed immunohistochemistry (IHC) (Tsujikawa et al., 2017) and CODEX (Goltsev et al., 2018) are *in situ* analyses of multiple marker channels over a large number of cells within a given tissue sample. These methods build upon dissociative single cell analysis methods like flow cytometry (Bradford et al., 2004) and single-cell RNA sequencing (Chen et al., 2019) to allow scientists to better understand spatial cell-cell interactions in biological samples.

One significant issue in multiplexed imaging data is the presence of systematic noise at a variety of levels, related to batch and slide effects, imaging variables, and optical effects (Berry et al., 2021; Chang et al., 2020). A single experiment may contain hundreds of slides and terabytes of data across which a researcher seeks to make inference (Maric et al., 2021). However, this data complexity and the within-slide dependencies induce complex effects that can disrupt inference. This technical variability can be compounded through the complex image pre-processing pipeline and may contribute to biases that increase type 1 or type 2 error. Further, it is difficult to develop a standardized pre-processing pipeline because of substantial variability in the markers used across different studies, as target proteins differ across organs and cancer types (Schapiro et al., 2021; Yapp et al., 2021).

Image normalization is a technique used to adjust the input pixel- or image-level values of an image to remove noise and improve image quality. Due to the nascent development of multiplexed imaging, there are few established statistical tools that address challenges related to technical variation in this data set (Chang et al., 2020). Normalization methods may improve similarity across images by removing the unknown effect of technical variability. Moreover, statistical methods for batch correction and image normalization can be modified to fit this complex data structure to ultimately reduce systematic noise and improve statistical inference.

Extensive work has been done in other fields to adjust for batch effects and systematic noise, particularly with regards to neuroimaging and genetic sequencing data. One primary method employed in both of these fields is the ComBat method, introduced for genetic micro-array data (Johnson et al., 2007) and then adapted to neuroimaging in the analysis of magnetic resonance imaging (MRI) data (Fortin et al., 2017; Yu et al., 2018). The ComBat method is a location-scale model that implements an empirical Bayes algorithm to adjust for batch effects, and is robust to outliers in small sample sizes. Curve registration, a non-parametric tool from functional data analysis, has been used in recent work to adjust for systematic variability in accelerometry and MRI data (Marron et al., 2015; Wrobel et al., 2020, 2019). In the neuroimaging context, curve registration is used to normalize the imaging data by non-linearly transform the image intensity domain so that it is similar across images from different subjects, potentially collected on different scanners.

While adaptable, existing methods for normalizing data from other domains cannot be directly applied within multiplexed imaging due to the unusual format of the data (cell populations can differ substantially across samples), and the heavy skewness of the image histogram. The few algorithms adapted specifically for normalizing multiplex imaging data still could benefit from upstream normalization using algorithms adapted from other domains (Chang et al., 2020; Raza et al., 2016). For example, the RESTORE algorithm is a method developed for multiplexed imaging that uses negative control cells to remove unwanted variation across slides (Chang et al., 2020). However, this method relies on clustering mutually exclusive marker pairs using cell-level labels that are defined using unnormalized marker intensities and thus embed biases as detailed in this paper. Raza et al also introduced normalization methods in the multiplexed domain that implement a procedure of image filters and transformations (Raza et al., 2016). These methods show improvements at the pixel and image level, but do not correct for slide or batch effects that are prevalent as detailed in this work. Hence, the normalization methods proposed here can be applied early in the image processing pipeline to reduce bias in subsequent steps like phenotyping and spatial correlation analyses.

In this paper we introduce and compare normalization and data transformation methods for multiplexed imaging data. These techniques combine transformations of the scale of the data from its raw form with algorithms (namely, ComBat and functional data registration) adapted to remove slide effects from the data. We further develop multiple novel metrics to quantify and measure the removal of technical variation in these data, where cell populations can differ across slides. We use data from the Human Tumor Atlas Network to evaluate the methods we compare here (Rozenblatt-Rosen et al., 2020). While we apply the methods here to segmented and quantified single-cell data from multiplexed imaging, they can also be applied at the pixel level.

## Methods

### Algorithm implementation

We compare 4 data transformations: log_10_, cube root, mean division (division by the slide-level mean), and mean division with log_10_, and 3 normalization procedures: no normalization, ComBat, and functional data registration, for a total of 12 potential multiplex image normalization algorithms (Table 1).

**Table 1:** Summary of normalization prodcedures implemented. Transformations (rows) and normalization (columns) performed on the data. Here *y* is the median cell intensity values for an arbitrary marker channel *c*, and *μ_ic_* is the slide mean for slide *i* of the median cell intensity values for marker channel *c*.

|  | Unnormalized | ComBat | Registration (fda) |
| --- | --- | --- | --- |
| $\log_{10}$ | $\log_{10}(y + 1)$ | ComBat ($\log_{10}(y + 1)$) | fda ($\log_{10}(y + 1)$) |
| Cube root | $\sqrt[3]{y}$ | ComBat ($\sqrt[3]{y}$) | fda ($\sqrt[3]{y}$) |
| Mean division | $\frac{y}{\mu_{ic}}$ | ComBat ($\frac{y}{\mu_{ic}}$) | fda ($\frac{y}{\mu_{ic}}$) |
| Mean division $\log_{10}$ | $\log_{10}(\frac{y}{\mu_{ic}} + \frac{1}{2})$ | ComBat ($\log_{10}(\frac{y}{\mu_{ic}} + \frac{1}{2})$) | fda ($\log_{10}(\frac{y}{\mu_{ic}} + \frac{1}{2})$) |

#### Transformations

Let *y* denote the raw data for a given marker channel, *c*. We consider the following transformations: the log_10_ transformation, log_10_(*y* + 1), where the addition of 1 follows since *y* is integer-valued; the cube root transformation, 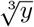; the mean division transformation: 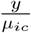, where *μ_ic_* is the mean intensity value on slide *i* for channel *c*; and the mean division log_10_ transformation, 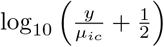, where again *μ_ic_* is the mean intensity value on slide *i* for channel *c*. Here the data are no longer integer-valued, and the addition of 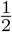 ensures values greater than 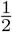 are positive and less than 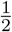 are negative to properly adjust this scale of data.

#### ComBat normalization

We adapted the empirical Bayes framework of the ComBat algorithm (Fortin et al., 2017; Johnson et al., 2007) for multiplexed imaging data. We parameterize mean and variance of the slide-level batch effects, with the location-scale model

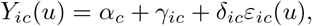

where we define *Y_ic_*(*u*) as the intensity of unit *u* on slide *i* for marker channel *c* and *α_c_* as the the grand mean of *Y_ic_*(*u*) for channel *c*. Though in principle units can be at the pixel or cell level, in our application, *Y_ic_*(*u*) is the median cell intensity (or its transformed counterpart) of a selected marker for a given segmented cell on a specific slide in the dataset. Here *γ_ic_* is the the mean batch effect of slide *i* for channel *c* and assume 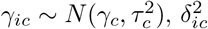 is the variance batch effect of slide *i* for channel *c* and assume 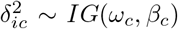, and we assume the random errors *ε_ic_*(*u*) ~ *N*(0,1). We use the data to estimate 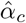 and then estimate 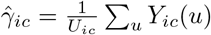, or the sample mean intensity on slide *i* for channel *c*. We further define 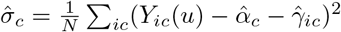 and let:

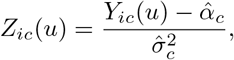

where we assume 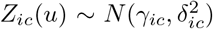. Based on the posterior conditional means, we find the following empirical Bayes estimators of the two batch effect parameters (a detailed derivation of these estimators can be found in the Supplement):

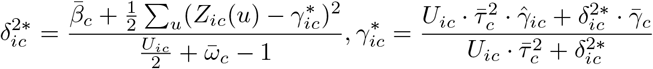

Where we define 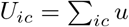, or the number of quantified cells present on a particular slide *i* for a given channel *c*. We calculate the hyper-parameter estimates of 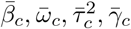 using the method of moments and iterate between estimating the hyper-parameters and batch effect parameters until convergence (Dempster et al., 1977). Upon convergence, we use these batch effects to adjust the data,

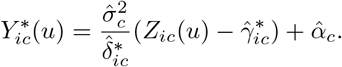

This model adjusts the Z-normalized intensity data, *Z_ic_*(*u*), by the mean and variance batch effects, and re-scales back to the initial scale of the data with the mean and variance of the raw marker intensity values. Note that zeroes were left in the data prior to the ComBat normalization, since for each scale transformation we perform on the data the zeroes are meaningful rather than an absence of signal.

#### Functional data registration

For the second normalization algorithm we implemented functional data registration using the `fda` R package (Ramsay and Silverman, 2005; Ramsay et al., 2020). This approach uses functional data analysis (FDA) methods to approximate the histograms for each slide and channel as smooth densities, and uses functional registration to align the densities to their average at the slide-level. Functional registration is performed by estimating a monotonic warping function for each density that stretches and compresses the intensities such that densities are aligned. These warping functions are then used to transform the marker intensity values in the images so that non-biological variability is reduced across slides.

Here, let our observed cell intensity values *Y_ic_*(*u*) have density *Y_ic_*(*u*) ~ *f*(*y* | *i*, *c*). Our goal is to remove technical variation related to the slide by estimating a warping function, *ϕ_ic_*(*y*), which is a monotonic transformation of the intensities. We first use a 21 degree of freedom cubic B-spline basis to approximate the densities of the median cell intensities for each slide and marker, *f*(*y* | *i*, *c*) ≈ *β^T^g*(*y*) where 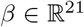 is an unknown coefficient vector and *g*(*y*) is a vector of known basis functions. We then register the approximated histograms to the average, restricting the warping function to be a 2 degree of freedom linear B-spline basis for some unconstrained functions *h*_1_(*y*) and *h*_2_(*y*) and for constants *C*_0_ and *C*_1_ to be estimated from the data,

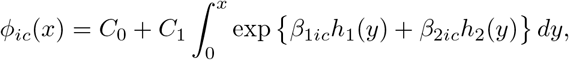

such that the transformation is monotonic (Ramsay and Silverman, 2005). Unknown parameters *β*_1*ic*_ and *β*_2*ic*_ are estimated to minimize,

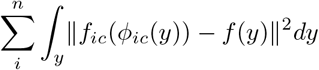

We then use *ϕ_ic_*(*y*) to calculate the normalized intensity values, 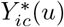:

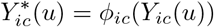

Note that the warping function *ϕ_ic_*(*y*) is a map that takes in the raw median cell intensity value and outputs a new, normalized intensity value. Images are then normalized by taking the original intensity values in the image, and transforming them using the map defined by the warping function. This combined process can be summarized as first taking the raw data, smoothing the histogram of these data using a B-spline basis expansion, and then calculating a warping function to transform the smoothed data so that densities across slides within marker channel *c* are aligned.

### Evaluation framework

There is no accepted gold standard for evaluating normalization methods in multiplexed imaging because the same tissue sample cannot be imaged precisely twice and there is substantial heterogeneity across samples (Nadarajan et al., 2019; Rozenblatt-Rosen et al., 2020). Here, our evaluation framework relies on the two following conditions to be deemed successful: (1) reduction in slide-to-slide variance in the cell intensity data and (2) preservation (and potential improvement) of existing biological signal in the data.

#### Visual alignment of marker densities

To determine if between-slide noise is visible when comparing densities, we visually inspect the changes in density curves for each transformation method. *A priori*, we expect that a successful transformation method will align the density curves across slides, and subsequently we inspect the placement of slide-level Otsu thresholds, a commonly used thresholding algorithm used in imaging analysis (Otsu, 1979), to confirm a reduction in variability between slides.

#### Otsu misclassification and accuracy

Otsu thresholding is a commonly used thresholding algorithm that defines an optimal threshold in grayscale images and histograms, maximizing the between-class variance of pixel values to separate the data into two classes (Otsu, 1979). In this use case, we define Otsu thresholds at the slide-level for each of the markers in the study, where a cell with intensity value greater than the Otsu threshold is deemed marker positive. We then compare this to a global Otsu threshold, combining all slides, for each marker to calculate a mean misclassification error across all slides for a given marker. For some marker channel *c*, slide *i*, and set of marker intensity values *y*, let *O_ic_*(*y*) be the values of *y* greater than the Otsu threshold for slide *i*, and let *O_c_*(*y*) be the values of *y* greater than the Otsu threshold across all slides *i* = 1, …, *N*. The misclassification metric is then defined as:

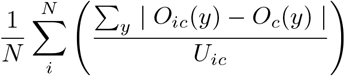

Here we calculate a slide-level misclassification error, e.g. the proportion of cells misclassified on each slide, and take an average of the misclassification error across slides for each marker channel. This measures the slide-to-slide agreement across all markers and transformation methods, to determine how similar Otsu thresholds are across slides following transformation.

We further implemented Otsu thresholding to compare definitions of a marker positive cell, with the test case using Otsu thresholding across slides to define a cell as marker positive and compared to the manual labels of CD3 and CD8 as marker positive cells (see the Dataset Section). This metric quantifies the accuracy of the Otsu thresholding method in recapitulating the bronze standard labels for each transformation method.

#### Proportions of variance

To further assess the removal of slide related variance following each transformation of the data, we fit a random effects model using the `lme4` R package (Bates et al., 2015) with a random intercept for slide to assess what proportion of variance in two markers, CD3 and CD8, is present at the slide-level. A successful normalization algorithm will reduce the slide-level variance, ultimately removing technical variability to improve the quality of the data.

#### Preservation of cell proportions

To measure the preservation of biological signal for each transformation method, we first quantified cell proportions in various tissue classes using Otsu thresholds. This method uses the manual labels for CD3- and CD8-positive cells to calculate the proportion of positive cells within each level of the data (e.g. slide identifier, slide region). This metric visualizes the change in baseline cell proportions of CD3- and CD8-positive cells for each transformation algorithm implemented using raincloud plots (Allen et al., 2019) to compare the distribution, box plot, and densities of marker positive cells.

#### UMAP embedding

The Uniform Manifold Approximation and Projection (UMAP) is a technique for dimension reduction (McInnes et al., 2018) commonly used in the biological sciences to distinguish differences in cell populations between single-cell data (Becht et al., 2019). Here we reduce the data into 2 UMAP embeddings for each of the transformation methods using only 4 markers in the dataset: vimentin, collagen, pancytokeratin, and Na^+^/K^+^-ATPase. These markers were chosen for their ability to easily distinguish epithelial and stromal cells. We expect the UMAP embeddings to yield clear separation of the data when using the epithelium label in our dataset (see the Dataset Section). Note that across each slide in the dataset, approximately 10% of the data was used to derive the UMAP embeddings to reduce computational and visualization time.

### Dataset

The data was collected from human colorectal cancer tissue samples from the Human Tumor Atlas Network (Rozenblatt-Rosen et al., 2020). The final dataset comprises over 2.2 million cells in the MxIF modality across over 2400 images on 43 different slides, with single-cell segmentation performed using an algorithm developed in-house (McKinley et al., 2019). Cell intensities for each marker were quantified as the median pixel value within the segmented cell, with tissue samples stained for 33 different marker channels. For the purpose of evaluating the algorithms compared in the paper, we restricted our attention to the following markers: beta catenin (BCATENIN), CD3D (CD3), CD8 (CD8), collagen (COLLAGEN), Na^+^/K^+^-ATPase (NAKATPASE), olfactomedin 4 (OLFM4), pan-cytokeratin (PANCK), SRY-Box 9 (SOX9), vimentin (VIMENTIN). These markers were chosen because of their ability to distinguish between epithelial and stromal cells, PANCK, COLLAGEN, NAKATPASE, VIMENTIN (Blom et al., 2017; Ijsselsteijn et al., 2019); as immune markers, CD3, CD8 (Galon et al., 2006); as stem cell markers, OLFM4, SOX9 (Van der Flier et al., 2009; Scott et al., 2010); and as implicated in colon cancer, BCATENIN, (Shang et al., 2017).

We used epithelial and stromal cell labels and manually labeled marker positive cells as biological variables in order to quantify loss or improvement of biological signal due to each normalization method. The epithelial labels were created for each slide at the image level using a random forest trained on all of the markers included in the dataset (for a complete list, see the Supplement). A cell was labeled as being in a particular cell class if that was the most likely class probability within the segmented cell area. We defined marker positive cells by first manually thresholding the immune marker images to create marker positive image masks. Then, for each segmented cell, the cell was defined as marker positive if more that 30% of its area contained marker pixels. We refer to these as manual labels for CD3 and CD8. We also used a tumor image mask to denote whether a cell is in a tumor-containing region.

## Results

### Removal of slide-to-slide variation

#### Visual alignment of marker densities

Density curves of the marker vimentin for each transformation algorithm and corresponding slide-level Otsu thresholds were compared to determine alignment of curves across slides after transformation (Figure 1). Beginning with the unnormalized values, the log_10_ and cube root methods produce density curves that are not well-aligned, contrasting the mean division and mean division log_10_ methods that both compress the scale of the data and align well across slides. Further, each ComBat method performs poorly at aligning and reducing noise in the data. This is likely due to the Gaussian assumptions of the ComBat model that are not met in either the bi-modal (log_10_, cube root, mean division log_10_) or right-skewed (mean division) methods. The functional data registration visually aligns the log_10_ and mean division log_10_ well, but does not perform as well with the cube root or mean division, potentially due to longer-tailed distributions of data that are not easily captured by the B-spline basis approximation.

**Figure 1:**
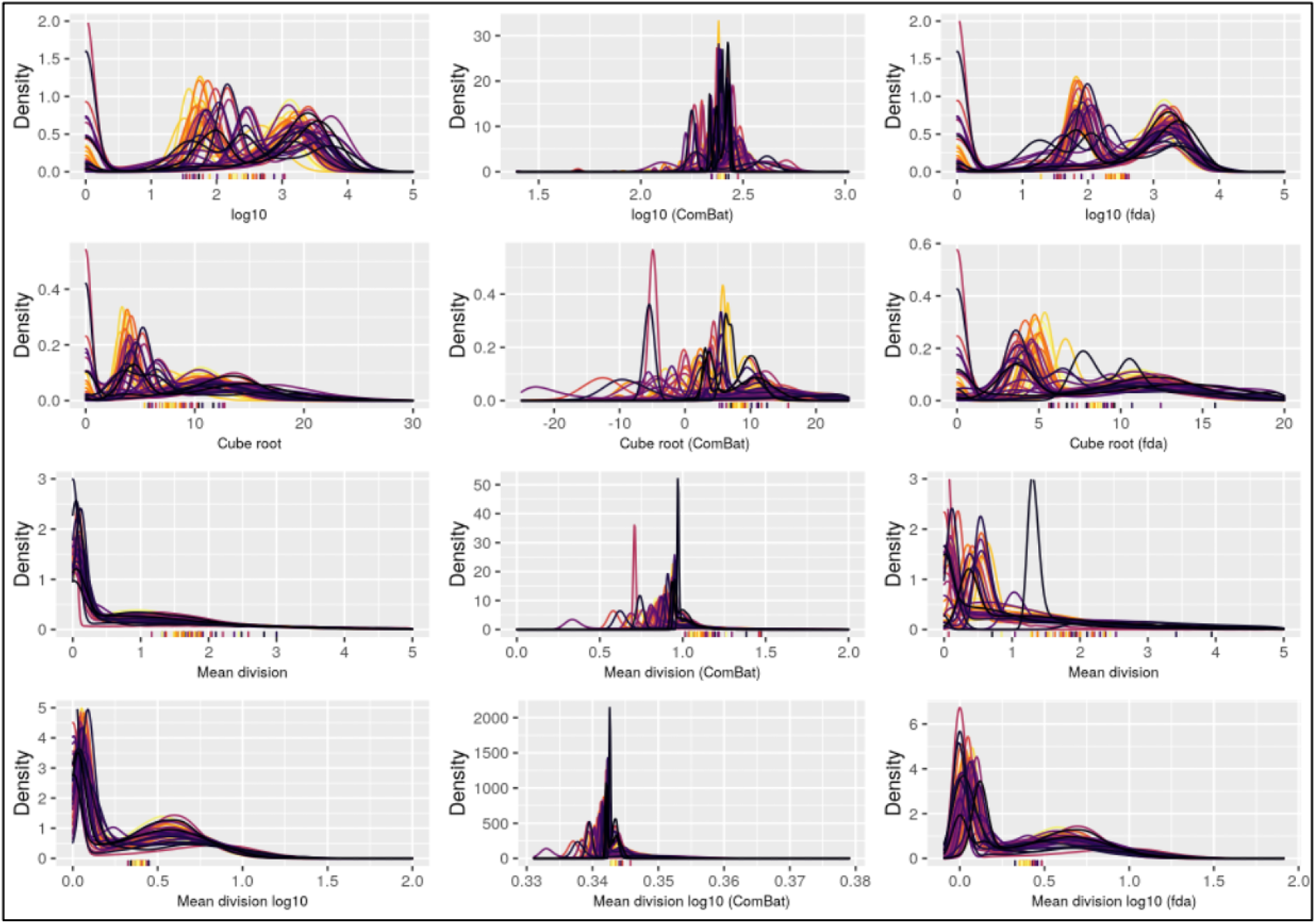
Visual comparison of vimentin marker densities for each transformation method. Density plots for the median cell intensity of the marker vimentin, where each color represents a different slide in the dataset. Each row is aligned with the scale transformations present in **Table 1**, where each column also matches with the normalization algorithms in **Table 1**. The ticks on the x-axis represent the Otsu thresholds for each slide for that transformed data, where the color again corresponds to the slide (such that the colors are one-to-one between threshold and density plot).

The best performing methods for this metric are the mean division log_10_ and mean division log_10_ combined with the functional data registration algorithm: the data is well-aligned across slides and the slide-level Otsu thresholds exhibit the least slide-to-slide variability. We also compared density curves of the markers CD3 and CD8 for each transformation algorithm, which largely present the same results (Supplementary Figures 1 and 2).

#### Otsu misclassification rate

In order to quantify how the normalization methods impact cell classification, we compared Otsu thresholding estimated at the slide level and across slides for each method to generate a misclassification rate and compare this to raw data (Figure 2A). Compared to the epithelium/stromal markers in the dataset, less identifiable markers like CD3 and CD8 yield the worst performance across nearly all methods, with large increases in the misclassification rate. Most methods increase the mean misclassification error relative to the unadjusted data, with the exception of the mean division, mean division log_10_, and the mean division log_10_ with functional data registration. This evaluation again aligns with earlier assessments and suggests that these methods present improvements in the slide-to-slide agreement across all markers compared to the unadjusted data.

**Figure 2:**
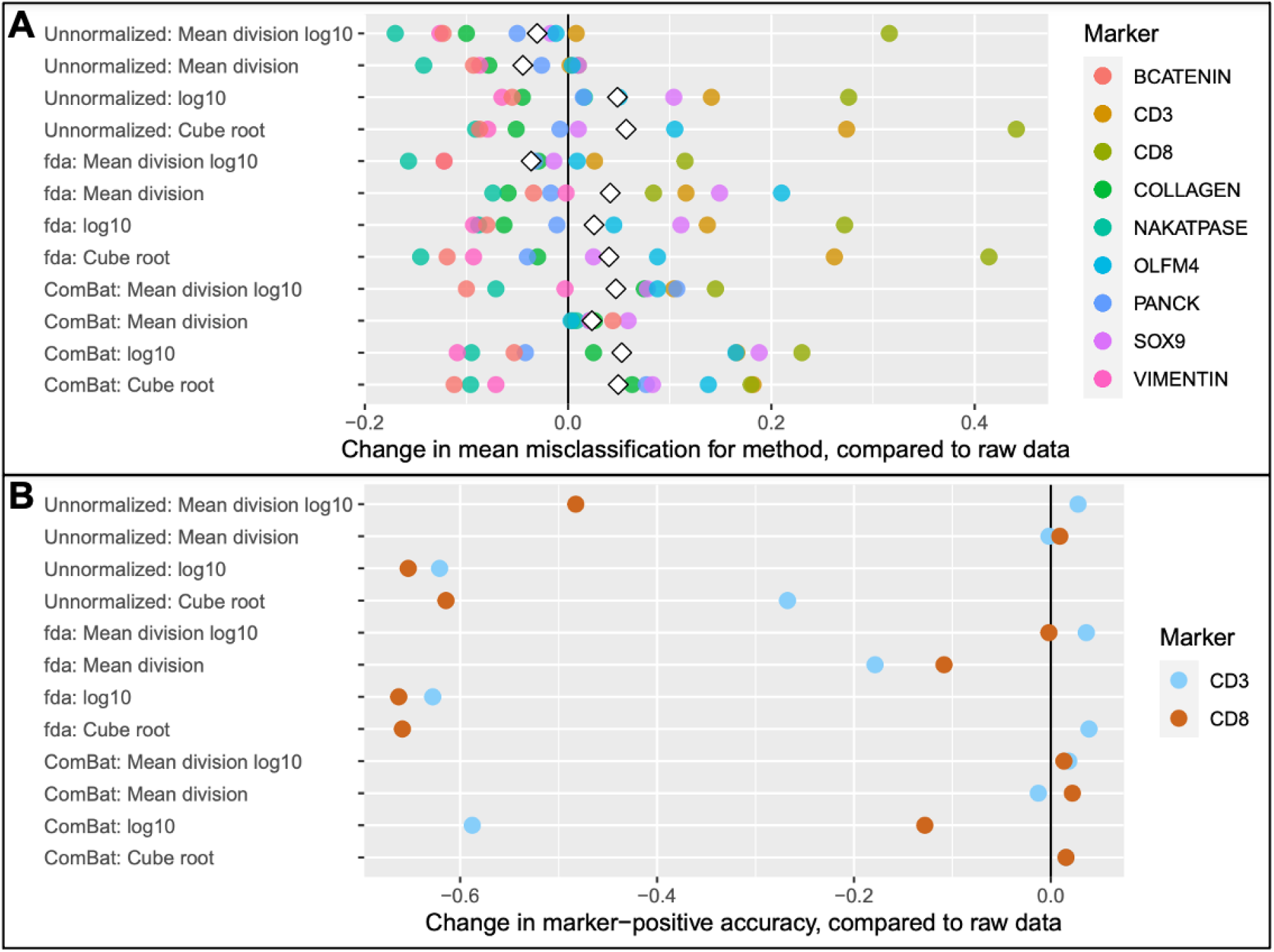
Otsu misclassification. (A) Otsu thresholds were calculated at the slide-level for each marker and compared to a global Otsu threshold for each marker to calculate a misclassification rate to compare transformation methods. The mean difference of the slide-level Otsu thresholds and the global Otsu thresold is then calculated for each marker, and the difference of this mean misclassification rate with the mean misclassification rate of the raw data is presented as a point for each of the 9 markers, with the white diamond representing the mean change in misclassification across all markers for a given method compared to the raw, unadjusted data. **Negative values indicate a reduction in misclassification error**. (B) Otsu thresholds were calculated across slides for each marker to determine marker positive cells, which were then compared to the manual labels for the markers CD3 and CD8 to determine the accuracy of defining a cell as marker positive. This is presented in the figure as a change in the accuracy of defining a cell as marker positive, compared to the accuracy in the raw data. **Positive values indicate an improvement in accuracy**.

#### Proportions of variance

To understand how well each method removes slide-related variability, we fit a random effects model on the median cell intensities after applying each combination of transformation and normalization. The ComBat algorithm, by design, removed all of the variability related to slide across all methods (Figure 3). Several of the algorithms increased variability related to slide (Figure 3; log_10_, cube root, mean division with registration, log_10_ with registration, cube root with registration). Unnormalized mean division, mean division log_10_, and mean division log_10_ with functional data registration reduced slide related variability and constitute an improvement according to this metric. While ComBat reduces slide variability, it completely removes slide effects that may include biological differences. The results of this metric suggest the potential utility of ComBat, and align with the first evaluation in terms of the ability of unnormalized mean division, mean division log_10_, and mean division log_10_ with functional data registration to reduce slide effects.

**Figure 3:**
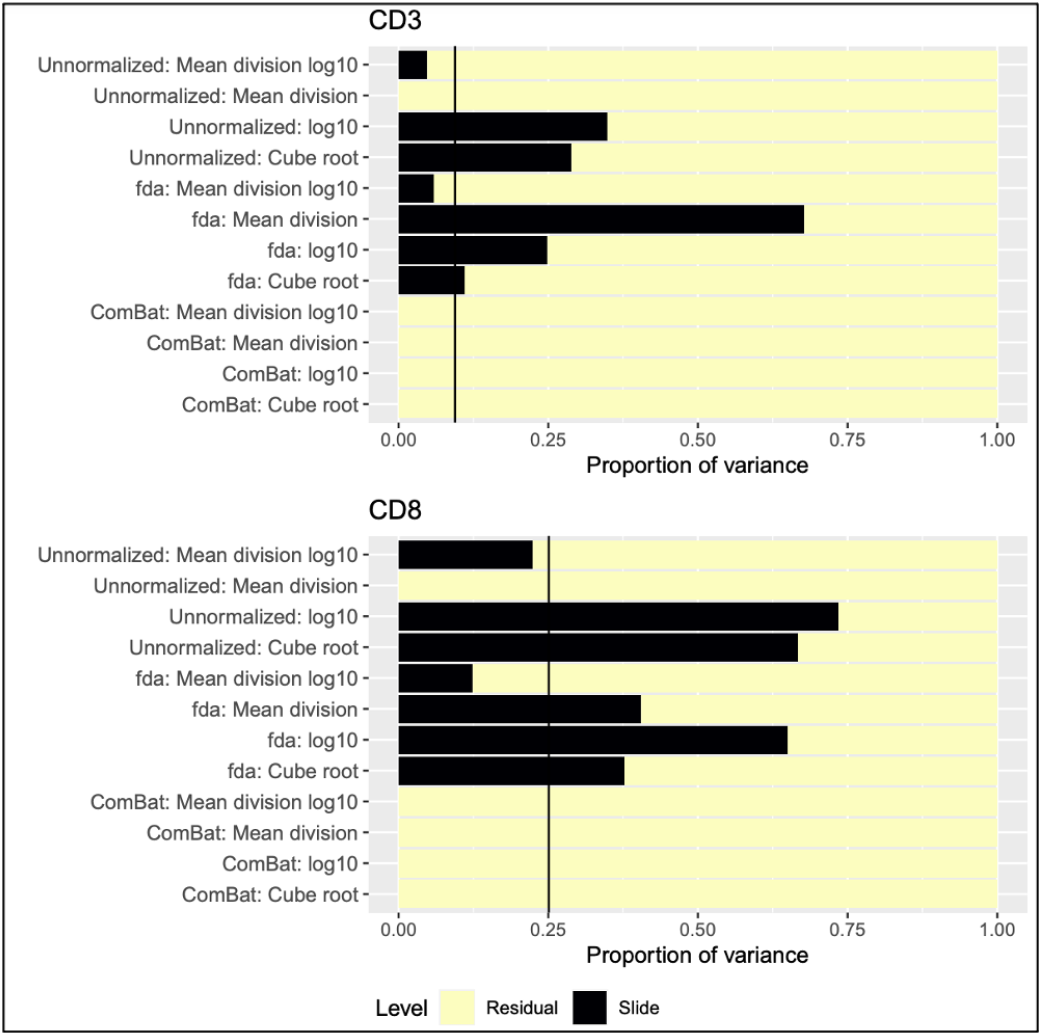
Proportion of variance present at slide-level in random effects model for CD3 and CD8 markers. Stacked bar charts that denote the proportion of variance at the slide-level (black) and residual variance (yellow) for each transformation method for the CD3 and CD8 markers. The vertical lines on each plot represent the proportion of variance at the slide-level in the raw, unadjusted data. Variance proportions were calculated using a random effects model with a random intercept for slide. Methods that perform well should reduce the slide level variance.

### Preservation of existing biological signal

#### Marker-positive accuracy using Otsu thresholds

We further utilized Otsu thresholding to identify marker positive cells and compared these to the manual labels for CD3 and CD8 to determine which normalization methods most accurately recapitulate the raw data (Figure 2B). Results suggest that the scale of the data is pivotal in whether a method maintains marker-positive accuracy, with each of the methods on the log_10_ scale demonstrating dramatic reductions in marker-positive accuracy compared to the raw data, while the mean division and mean division log_10_ methods perform the best across all methods (excluding a reduction in CD8 accuracy for unnormalized mean division log_10_). The methods that performed well in the aforementioned evaluation metrics perform well here, namely the mean division method and the mean division log_10_ with functional data registration. This continues to suggest these methods reduce the slide-to-slide variation present in the data while accurately capturing marker-positive cells after transformation.

#### Preservation of cell proportions

We compared CD3 and CD8 cell proportions within epithelium and stroma using the proportions estimated by Otsu thresholding after each normalization method to quantify the preservation of biological signal for each method, as compared to those estimated using the manual labels (Figure 4). For both CD3 and CD8, we see that the log_10_ scale does not replicate the cell proportions from the manual labels, while the mean division log_10_ performs well in both markers across normalization algorithms (again excluding the unnormalized mean division log_10_ for CD8). Per this metric in both CD3 and CD8, and re-affirming prior evaluation, the mean division method and the mean division log_10_ with functional data registration maintain the cell proportions most closely. This again points to the ability of these methods to robustly maintain biological signal in the unadjusted data while removing slide-to-slide variation.

**Figure 4:**
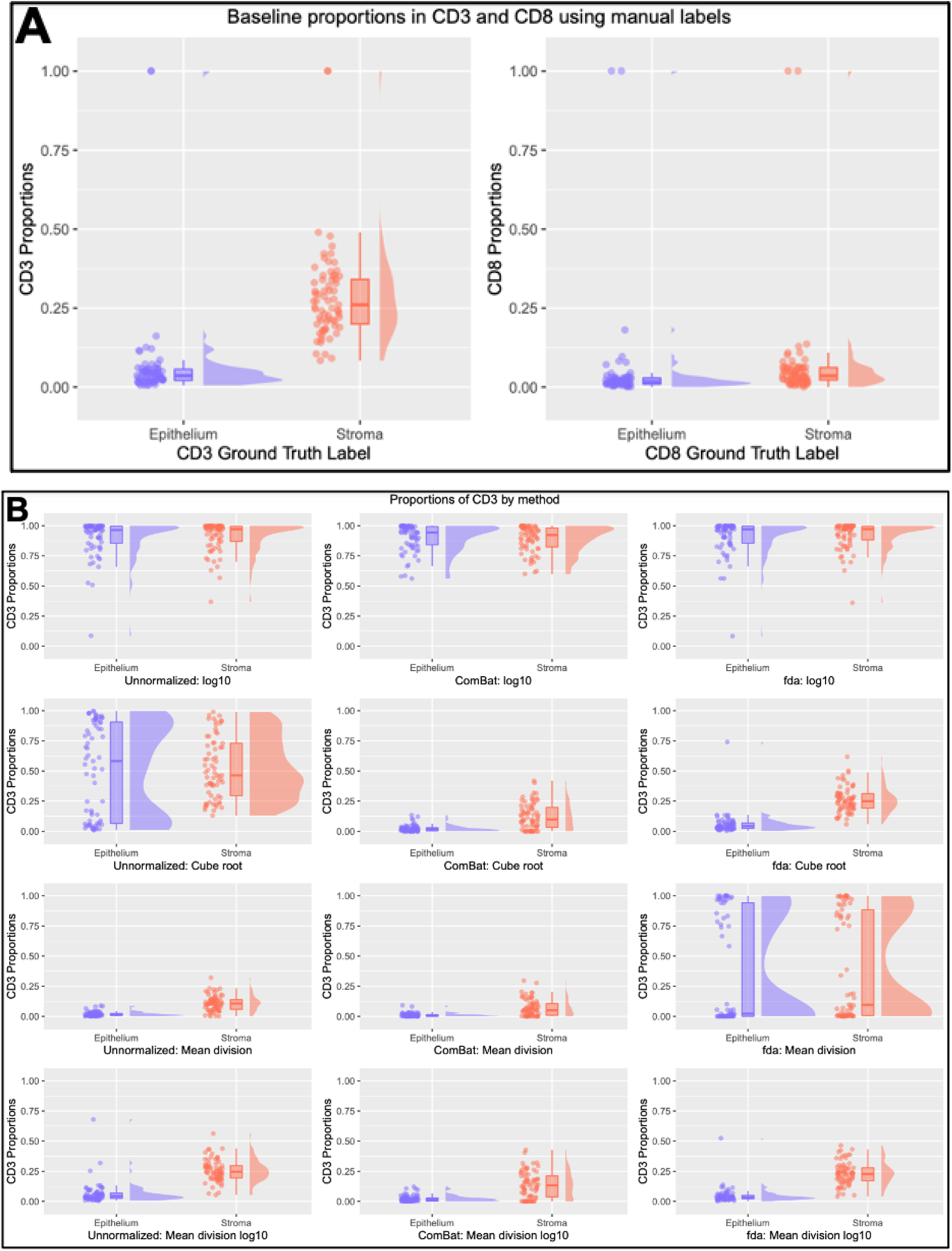
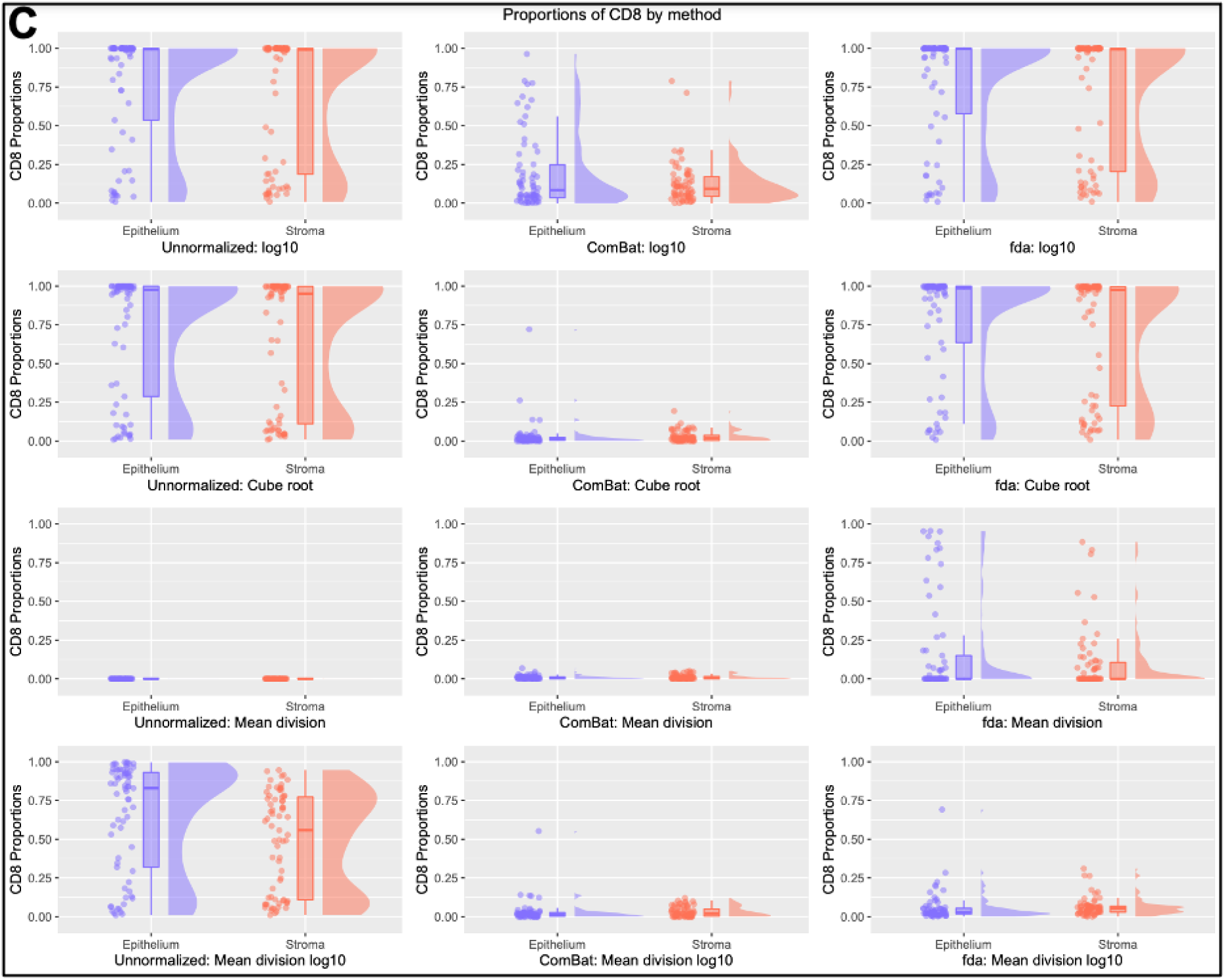
Comparison of cell proportions for each transformation method. A comparison of estimated proportions for the manually labeled cells for CD3 and CD8 (A) to the proportions of (B) CD3 and (C) CD8 for each normalization methods. Positive cells for the normalization methods are determined by Otsu thresholding across all slides. Methods that maintain similar estimates to the manual labels are considered more accurate.

#### UMAP embedding

We compared UMAP embeddings of four related markers across normalization methods to compare the separation of epithelium and stromal tissue labels. In the raw data, the embeddings separate well, however the data includes the presence of outliers that suggest mixing of the tissue classes in the UMAP embedding space (Figure 5A). Results on the log_10_ and cube root scales display co-localization and further clustering of results that does not clearly depict separation as desired (Figure 5C). We do observe distinct separation of the aforementioned methods of interest: mean division, mean division log_10_, and the mean division log_10_ with functional data registration - each of these UMAP embeddings presents distinct groups divided along a single plane that suggests these methods are improving the separation of these two tissue classes.

**Figure 5:**
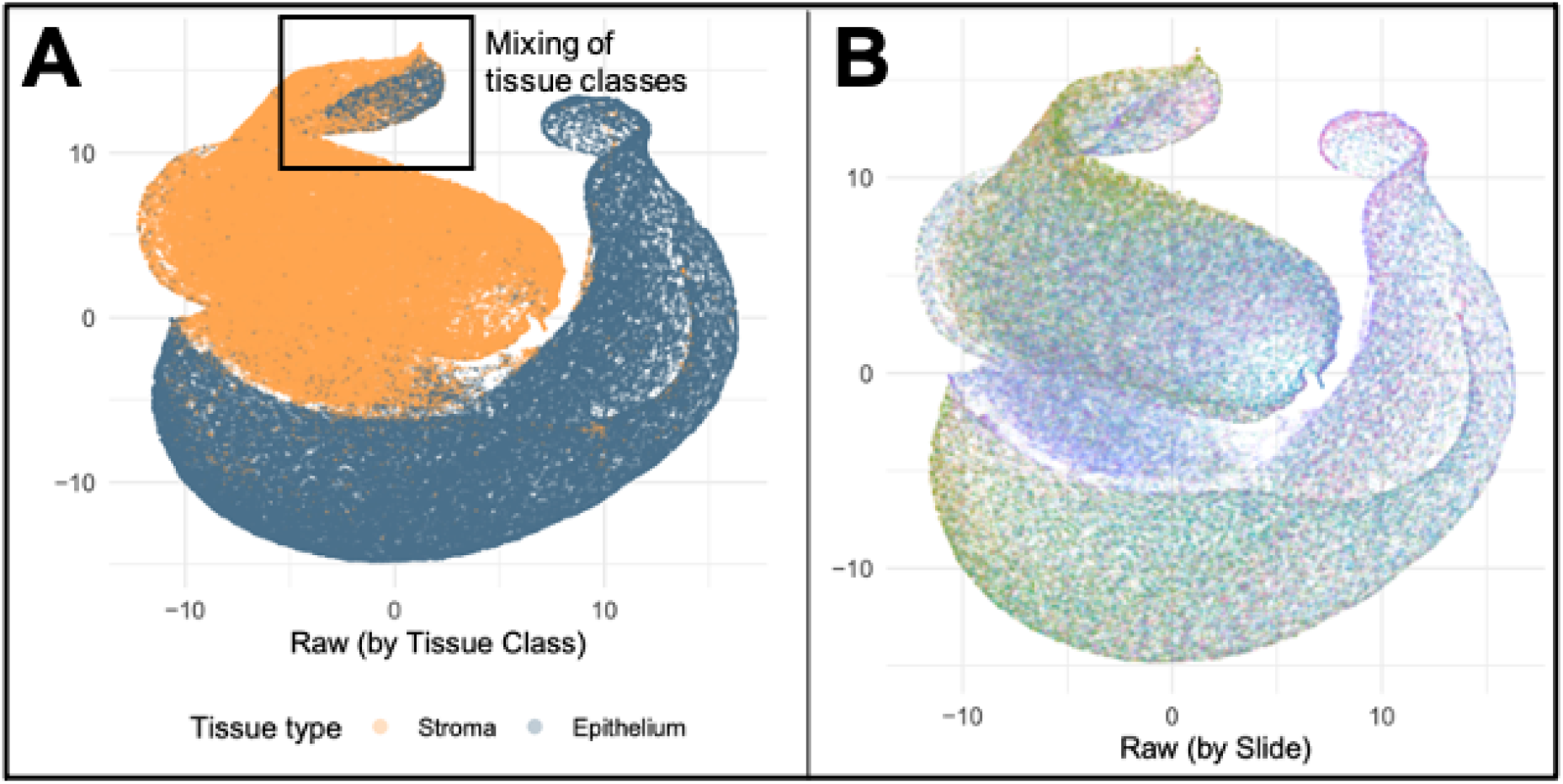
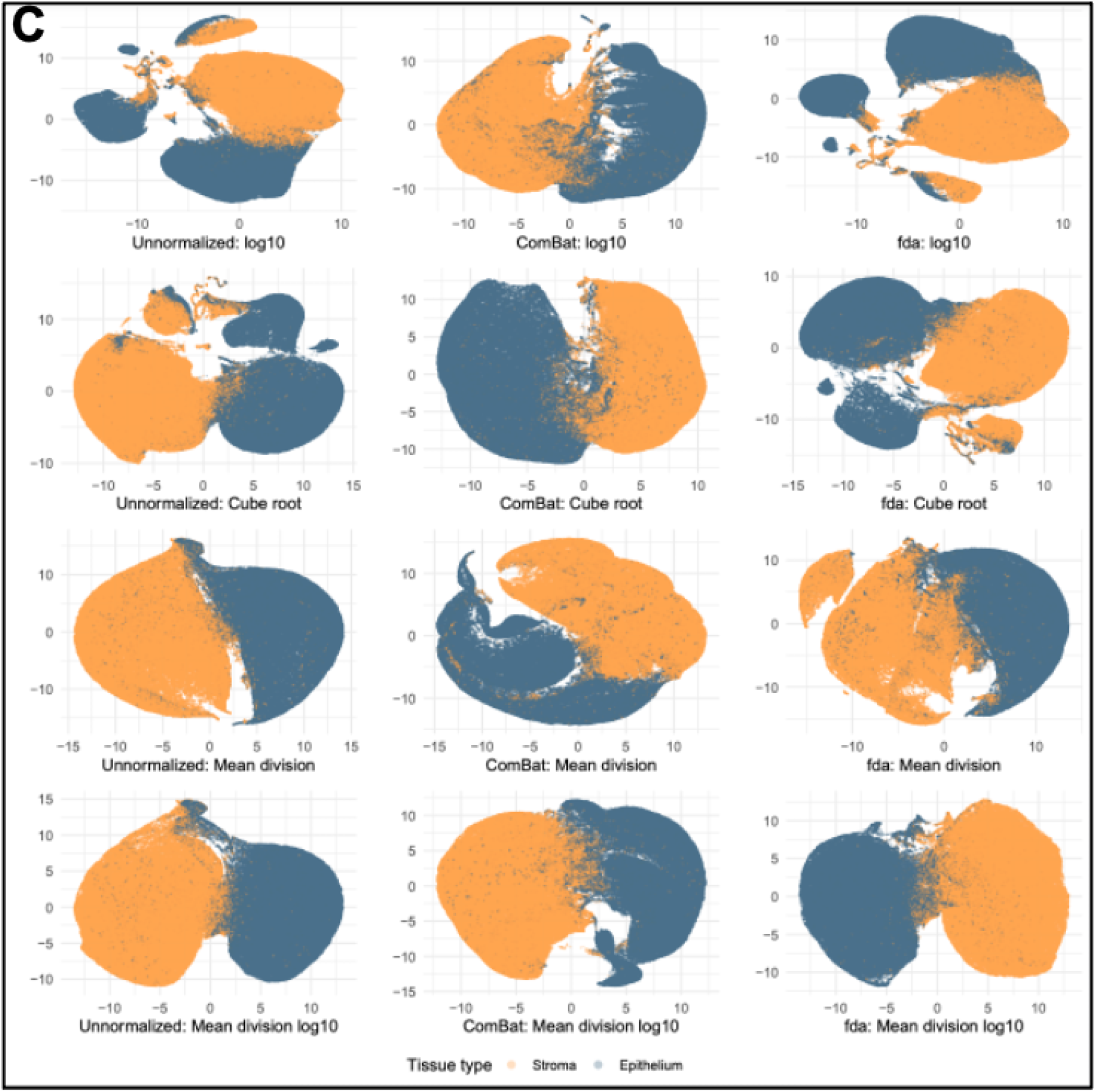
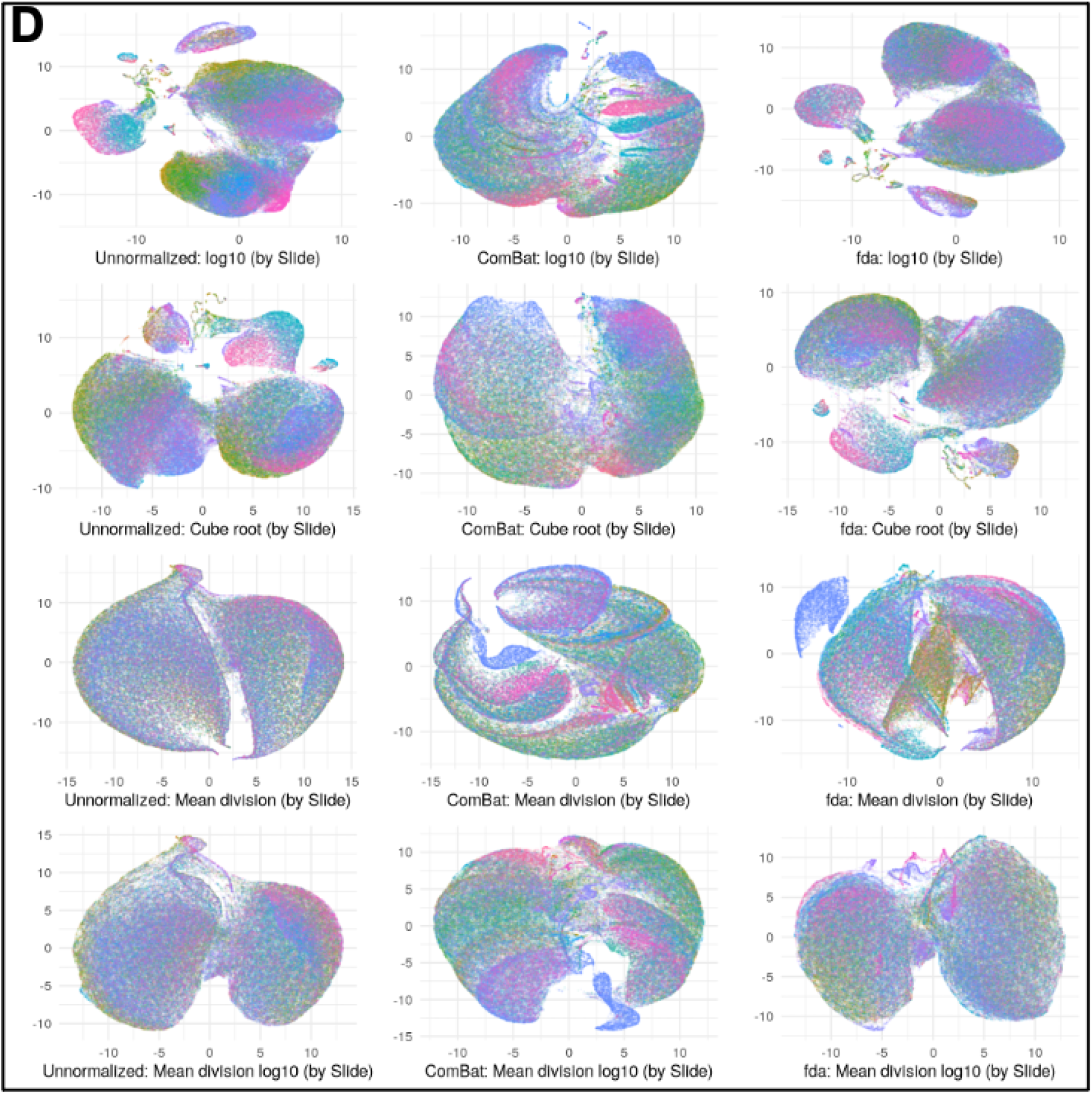
UMAP embedding of data for each transformation method. UMAP embedding of the raw, unadjusted data with points colored by tissue type (A) and slide identifier (B), compared to UMAP embeddings for each transformation method in the study with points colored by tissue type (C) and slide identifier (D). The rectangle in (A) denotes the mixing of tissue classes present in the raw, unadjusted data UMAP embedding.

We also compared the distribution of the unique slide identifiers in the UMAP embeddings of these four markers, which in the raw data points to specific slide co-localization in the data (Figure 5B). Normalization methods like the unnormalized log_10_ and cube root transformations, as well as each of the ComBat normalized data, worsen the distribution of these slide identifiers and present clear slide effects present in the adjusted data as demonstrated by regions largely composed of the same color (Figure 5D). This suggests that ComBat removes both biological signal and slide-to-slide effects that are exaggerated in the UMAP embedding space. In contrast, there is reduced slide-to-slide clustering in the UMAP embeddings for each of the following methods: mean division, mean division log_10_, and mean division log_10_ with functional data registration. These methods appear to both reduce the observed slide-to-slide variation noted here and in the aforementioned results, while maintaining necessary biological signal of interest.

## Discussion

In this paper, we derived the ComBat algorithm for a new modality and employed a novel use of functional data registration to align histograms of multiplexed imaging data. In the absence of a gold standard for comparison in multiplexed imaging data, validating any normalization procedure is challenging. The suggested evaluation framework introduced here can be used to assess the presence and reduction of slide effects in multiplexed imaging data, which we implemented to evaluate 12 combinations of transformations and normalization methods. Further, our framework can be applied in the absence of a ground truth by quantifying the amount of slide related variability and comparing to manually labeled biological features, providing a foundation for further development of evaluation criteria in the multiplexed domain.

We find that the raw data scale has clear slide-to-slide variation present, and that normalization methods can reduce slide level variation while preserving and improving biological signal relative to the raw, unadjusted data. These findings suggest that the mean division transformation method reduces slide variability and improves the biological signal. In addition, the mean division log_10_ scale (unnormalized) performs well across all evaluation metrics, with the noted exclusion of results for the marker CD8. This discrepancy is remedied with the functional data registration, which is a limitation of the mean division log_10_ transformation but points to the robustness of the registration algorithm to maintain and improve the quality of the data.

However, note that the registration algorithm does not perform well with skewed data, suggesting that improvements we see in data that appears bi-modal (e.g., better suited to the non-parametric assumption of functional data) is not necessarily transferable to right-skewed data that violates assumptions of smoothness in the B-spline basis - future work could explore this result. The ComBat method performs adequately, but appears to over normalize the data and relies heavily on a Gaussian assumption that is violated in this skewed-right dataset. Recent adaptations of ComBat like ComBat-seq for RNA-seq data may provide a better framework to implement in the multiplexed imaging space (Zhang et al., 2020), including future work that could address how the algorithm handles zeroes.

In practice, the mean division method is simple, computationally efficient, and less likely to introduce error while still reducing slide-to-slide variation and maintaining biological signal. The mean division log_10_ method may be necessary in the case of statistical modeling, since skewed distributions are not suitable for many statistical models, but may not the best way to represent cell intensities as a predictor variable (as appears the case for the mean division method). We see that in the case of mean division log_10_ data, it may be necessary to use the registration algorithm to remedy discrepancies like those visible for the marker CD8.

Similarly, the use of Otsu thresholding in this paper is typical for imaging domains, but future work may suggest that a separately optimized thresholding algorithm may yield superior results. Notably, the correspondence between an marker positive cell defined by an Otsu threshold and biological signal is not necessarily one-to-one. For example, the log_10_ transformation non-linearly compresses the domain, such that a larger proportion of the x-axis is allotted to cells that are marker negative (background and unexpressed cells), which may have led to greater variability in the Otsu thresholds.

## Funding

This work was supported by the National Institutes of Health (T32LM012412 to C.H., R01DK103831 and U01CA215798 to K.L., U2CCA233291 to R.C., K.L., M.S.), the Colorado Clinical and Translational Sciences Institute (UL1TR002535) and the Vanderbilt Ingram Cancer Center GI SPORE (P50CA236733). Study activities were conducted in part by the Survey and Biospecimen Shared Resource (P30CA68485), the Tissue Pathology Shared Resource (P30CA068485, U24DK059637), the Digital Histology Shared Resource, the NCI Cooperative Human Tissue Network (CHTN) Western Division (UM1CA183727), and REDCap (UL1TR000445).

## Notes

https://github.com/statimagcoll/MultiplexedNormalization

### Competing Interest Statement

The authors have declared no competing interest.

