## Supplemental Information for "Quantifying and correcting slide-to-slide variation in multiplexed immunofluorescence images"

### Supplementary Materials

#### Application of ComBat

Note from the **Methods** section of the paper that we have assumed that the standardized data $Z_{ic}(u) \sim N(\gamma_{ic}, \delta_{ic}^2)$ with the following priors on the batch effects:

$$\gamma_{ic} \sim N(\gamma_c, \tau_c^2), \delta_{ic}^2 \sim IG(\omega_c, \beta_c)$$

Recall that $i$ denotes the slide from which the data was collected, $c$ denotes the marker of interest, and $u$ defines the unit of measuring intensity, which for this study is the median quantified marker intensity of the segmented cell. Note also that we defined $U_{ic} = \sum_{ic} u$, or the number of quantified cells present on a particular slide $i$ for a given channel $c$.

##### Posterior Derivation for $\gamma_{ic}$

Using the empirical Bayes methodology, we must derive the posterior mean estimator of $\gamma_{ic}$ to utilize in the ComBat model. Hence:

$$\begin{aligned}
\pi\left(\gamma_{ic}|Z_{ic}(u), \delta_{ic}^2\right) &= L\left(Z_{ic}(u)|\gamma_{ic}, \delta_{ic}^2\right) \cdot \pi(\gamma_{ic}) \\
&\propto \exp\left\{-\frac{1}{2\delta_{ic}^2}\sum_u \left(Z_{ic}(u) - \gamma_{ic}\right)^2\right\} \cdot \exp\left\{-\frac{1}{2\tau_c^2}(\gamma_{ic} - \gamma_c)^2\right\} \\
&= \exp\left\{-\frac{1}{2\delta_{ic}^2}\left(\sum_u Y_{ic}^2(u) - 2\sum_u Z_{ic}(u)\gamma_{ic} + U_{ic} \cdot \gamma_{ic}^2\right) - \frac{1}{2\tau_c^2}\left(\gamma_{ic}^2 - 2\gamma_{ic}\gamma_c + \gamma_c^2\right)\right\} \\
&\propto \exp\left\{-\frac{1}{2}\left(\frac{U_{ic}\tau_c^2 + \delta_{ic}^2}{\delta_{ic}^2\tau_c^2}\right)\left[\gamma_{ic}^2 - 2\left(\frac{\tau_c^2\sum_u Z_{ic}(u) + \delta_{ic}^2\gamma_c}{U_{ic}\tau_c^2 + \delta_{ic}^2}\right)\gamma_{ic}\right]\right\}
\end{aligned}$$

Which after we complete the square, we see this is posterior follows the Normal distribution with the following expectation:

$$E\left[\gamma_{ic}|Z_{ic}(u), \delta_{ic}^2\right] = \frac{\tau_c^2\sum_u Z_{ic}(u) + \delta_{ic}^2\gamma_c}{U_{ic}\tau_c^2 + \delta_{ic}^2}$$

To derive an estimator of the batch effect parameter, we must define the following estimators of the hyperparameters:

$$\bar{\gamma}_c = \frac{1}{U_{ic}}\sum_i \hat{\gamma}_{ic} \text{ and, } \bar{\tau}_c^2 = \frac{1}{U_{ic} - 1}\sum_i (\hat{\gamma}_{ic} - \bar{\gamma}_c)^2$$

Hence we now derive the following estimator of $\gamma_{ic}$:

$$\gamma_{ic}^* = \frac{\bar{\tau}_c^2 U_{ic}\hat{\gamma}_{ic} + \delta_{ic}^{2*}\bar{\gamma}_c}{U_{ic}\bar{\tau}_c^2 + \delta_{ic}^{2*}}$$

**Posterior Derivation for $\delta_{ic}^2$**

We employ the same methodology to derive the posterior mean estimator of $\delta_{ic}^2$:

$$\begin{aligned}
\pi\left(\delta_{ic}^2 | Z_{ic}(u), \gamma_{ic}\right) &= L\left(Z_{ic}(u)|\gamma_{ic}, \delta_{ic}^2\right) \cdot \pi(\delta_{ic}^2) \\
&\propto {\delta_{ic}^2}^{-\frac{U_{ic}}{2}} \exp\left\{-\frac{1}{2\delta_{ic}^2}\sum_u \left(Z_{ic}(u)-\gamma_{ic}\right)^2\right\} \cdot {\delta_{ic}^2}^{-(\omega_c+1)} \exp\left\{-\frac{\beta_c}{\delta_{ic}^2}\right\} \\
&= {\delta_{ic}^2}^{-\left(\left[\frac{U_{ic}}{2}+\omega_c\right]+1\right)} \exp\left\{-\frac{1}{2\delta_{ic}^2}\left(\sum_u Y_{ic}^2(u) - 2\sum_u Z_{ic}(u)\gamma_{ic} + U_{ic}\cdot\gamma_{ic}^2\right) - \frac{\beta_c}{\delta_{ic}^2}\right\} \\
&\propto {\delta_{ic}^2}^{-\left(\left[\frac{U_{ic}}{2}+\omega_c\right]+1\right)} \exp\left\{-\frac{1}{\delta_{ijc}^2}\left(\beta_c + \frac{1}{2}\sum_u \left(Z_{ic}(u)-\gamma_{ic}\right)^2\right)\right\}
\end{aligned}$$

Which we note is an Inverse Gamma distribution with the following expectation:

$$E\left[\delta_{ic}^2 | Z_{ic}(u), \gamma_{ic}\right] = \frac{\beta_c + \frac{1}{2}\sum_u \left(Z_{ic}(u)-\gamma_{ic}\right)^2}{\frac{U_{ic}}{2}+\omega_c - 1}$$

To derive an estimator of the batch effect parameter, we must define the following estimators:

$$\hat{\delta}_{ic}^2 = \frac{1}{U_{ic}-1}\sum_u (Z_{ic}(u) - \hat{\gamma}_{ic})^2$$

We then calculate the sample mean of the $\hat{\delta}_{ic}^2$, $\bar{M}_c$ and $\bar{S}_c^2$ and set these equal to the moments of an Inverse Gamma distribution to yield the following estimators:

$$\bar{\omega}_c = \frac{\bar{M}_c + 2\bar{S}_c^2}{\bar{S}_c^2} \text{ and, } \bar{\beta}_c = \frac{\bar{M}_c^3 + \bar{M}_c\bar{S}_c^2}{\bar{S}_c^2}$$

Hence we now derive the following estimator of $\delta_{ic}^2$:

$$\delta_{ic}^{2*} = \frac{\bar{\beta}_c + \frac{1}{2}\sum_u \left(Z_{ic}(u)-\hat{\gamma}_{ic}\right)^2}{\frac{U_{ic}}{2}+\bar{\omega}_c - 1}$$

#### CD3 and CD8 Density Plots

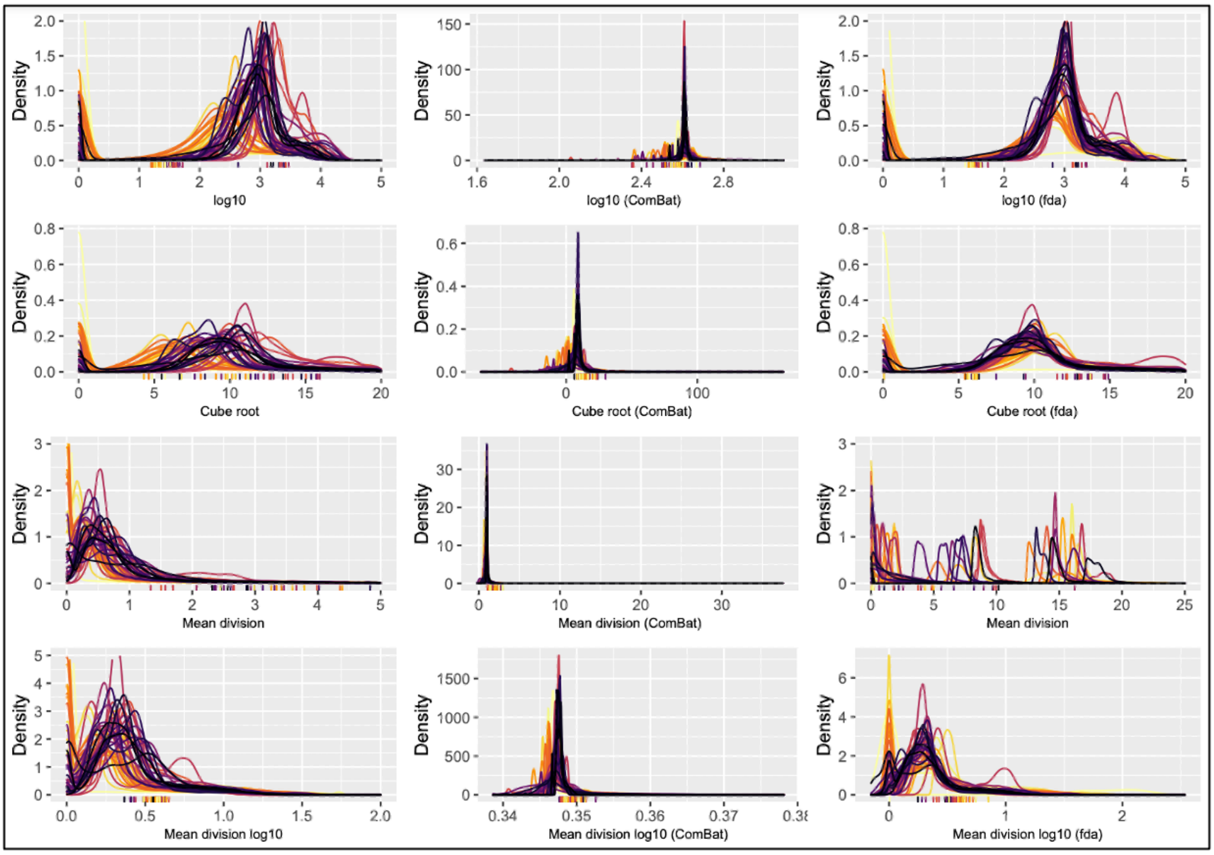


**Supplementary Figure 1: Visual comparison of CD3 marker densities for each transformation method**

Density plots for the median cell intensity of the marker CD3, where each color represents a different slide in the dataset. Each row is aligned with the scale transformations present in **Table 1**, where each column also matches with the normalization algorithms in **Table 1**. The ticks on the x-axis represent the Otsu thresholds for each slide for that transformed data, where the color again corresponds to the slide (such that the colors are one-to-one between threshold and density plot).

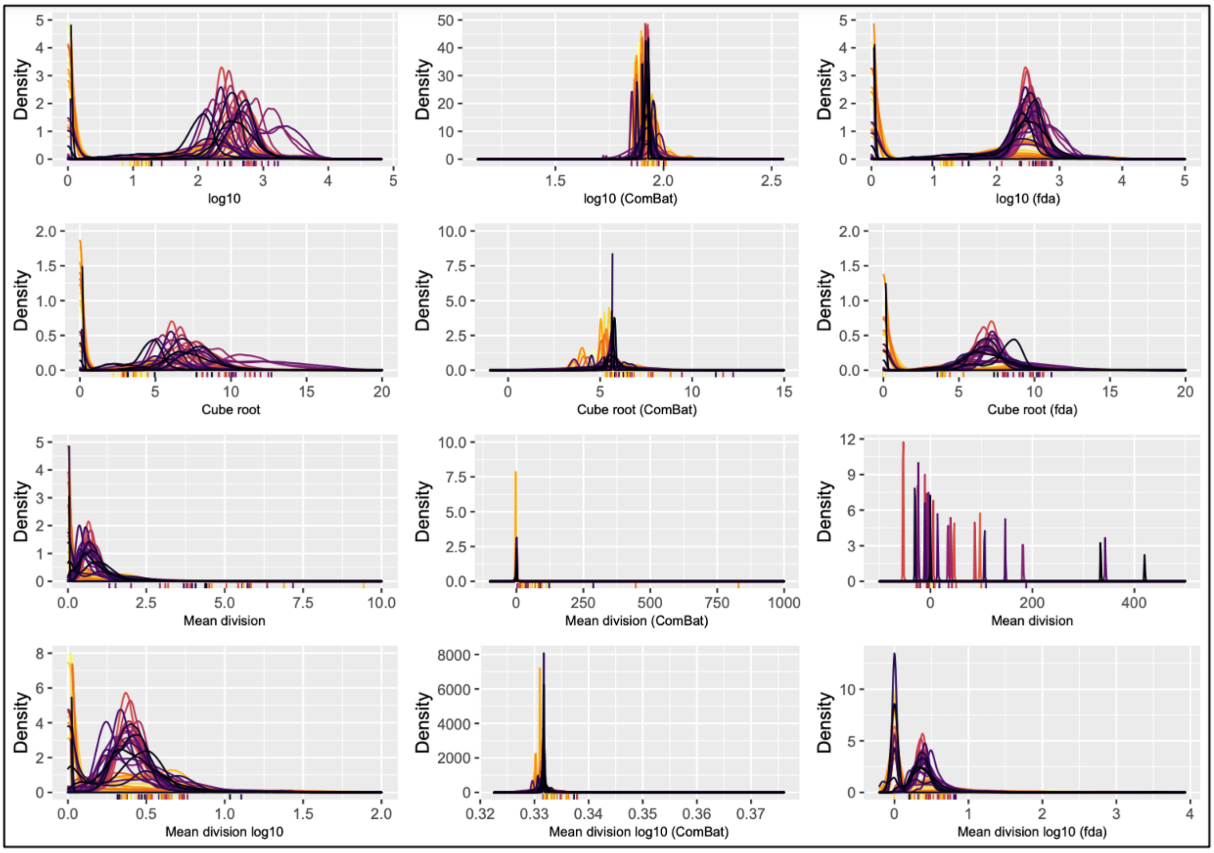


**Supplementary Figure 2: Visual comparison of CD8 marker densities for each transformation method**

Density plots for the median cell intensity of the marker CD8, where each color represents a different slide in the dataset. Each row is aligned with the scale transformations present in **Table 1**, where each column also matches with the normalization algorithms in **Table 1**. The ticks on the x-axis represent the Otsu thresholds for each slide for that transformed data, where the color again corresponds to the slide (such that the colors are one-to-one between threshold and density plot).

#### Marker List

| Dataset definition | Marker |
|---|---|
| ACTININ | Actinin |
| BCATENIN | Beta Catenin |
| CD11B | CD11b |
| CD20 | CD20 |
| CD3 | CD3D |
| CD4 | CD4 |
| CD45 | Protein tyrosine phosphatase, receptor type, C |
| CD68 | CD68 |
| CD8 | CD8 |
| CGA | Chromogranin A |
| COLLAGEN | Collagen |
| COX2 | Cyclooxygenase-II |
| ERBB2 | Erb-B2 Receptor Tyrosine Kinase 2 |
| FOXP3 | Forkhead Box P3 |
| HLAA | Human Leukocyte Antigen |
| LYSOZYME | Lysozyme |
| MUC2 | Mucin 2 |
| NAKATPASE | $Na^{+}/K^{+}$-ATPase |
| OLFM4 | Olfactomedin 4 |
| PANCK | Pan-Cytokeratin |
| PCNA | Proliferating Cell Nuclear Antigen |
| PEGFR | phospho-Epidermal Growth Factor Receptor |
| PSTAT3 | Phosphorylated Signal Transducer and Activator of Transcription |
| SMA | $\alpha$-Smooth Muscle Actin |
| SNA | Spherical Nucleic Acid |
| SOX9 | SRY-Box 9 |
| VIMENTIN | Vimentin |
| DAPI | $4'$,6-diamidino-2-phenylindole |
| CD45B | CD45RB |
| GACTIN | Globular Actin |
| PDL1 | Programmed Death-Ligand 1 |
| CDX2 | Caudal-type Homeobox 2 |
| MUC5AC | Mucin 5AC |
